# Dynamics of the sex ratio in *Tetrahymena thermophila*

**DOI:** 10.1101/327544

**Authors:** Guangying Wang, Kai Chen, Jing Zhang, Xuefeng Ma, Shanjun Deng, Jie Xiong, Xionglei He, Yunxin Fu, Wei Miao

## Abstract

Sex is often hailed as one of the major successes in evolution, and in sexual organisms the maintenance of proper sex ratio is crucial. As a large unicellular eukaryotic lineage, ciliates exhibit tremendous variation in mating systems, especially the number of sexes and the mechanism of sex determination (SD), and yet how the populations maintain proper sex ratio is poorly understood. Here *Tetrahymena thermophila*, a ciliate with seven mating types (sexes) and probabilistic SD mechanism, is analyzed from the standpoint of population genetics. It is found based on a newly developed population genetics model that there are plenty of opportunities for both the co-existence of all seven sexes and the fixation of a single sex, pending on several factors, including the strength of natural selection. To test the validity of predictions, five experimental populations of *T. thermophila* were maintained in the laboratory so that the factors that can influence the dynamics of sex ratio could be controlled and measured. Furthermore, whole-genome sequencing was employed to examine the impact of newly arisen mutations. Overall, it is found that the experimental observations highly support theoretical predictions. It is expected that the newly established theoretical framework is applicable in principle to other multi-sex organisms to bring more insight into the understanding of the maintenance of multiple sexes in a natural population.

## Introduction

Sex is often hailed as one of the most successful evolutionary inventions across eukaryotes (Weismann 1887; Bell 1982) and is fundamental for lineage survival (Speijer *et al.* 2015). This is due to various benefits provided by sex, such as speeding up adaptation by accelerating the accumulation of beneficial mutations (Fisher 1930; Muller 1932) and allowing natural selection to proceed more effectively by increasing the genetic variance in fitness (Weismann 1887; Burt 2000). For most sexual organisms, a crucial prerequisite for the occurrence of sex is that a population must maintain appropriate proportion of different sexes, or sex ratio. The strategies to adjust offspring sex ratio have been well demonstrated in a wide range of organisms with two sexes, male and female, in the context of Fisher’s equal allocation theory and its extensions (West 2009). However, how organisms with multiple (>2) sexes maintain proper sex ratio in the population is still poorly understood.

Ciliates are a large unicellular eukaryotic evolutionary lineage that show rapid diversification in many aspects of the mating systems (Phadke and Zufall 2009). First, ciliates exhibit a great variation in the number of mating types (sexes), ranging from two to several or more, e.g. 100 for *Stylonychia mytilus* (Ammermann 1982). Second, the mechanisms of sex determination (SD) differ widely, ranging from Mendelian systems to developmental nuclear differentiation, either stochastic or cytoplasmic (Orias *et al.* 2017). The well-studied ciliate, *Tetrahymena thermophila*, has seven self-incompatible sexes (I—VII) that are determined by alleles at a single locus (*mat*). Unlike the sex-specific alleles (*mat-a, mat-alpha*) in yeast, each *mat* allele specifies the probability with which a progeny cell will express one of the seven sexes (Arslanyolu and Doerder 2000). For example, a classic B-type *mat* allele specifies the following probabilities for each sex: I, 0; II, 0.275; III, 0.192; IV, 0.278; V, 0.076; VI, 0.041; VII, 0.138 (Nanney 1960). This particular form of sex inheritance has been called probabilistic SD and the unique distribution of probabilities is called the allele’s SD pattern (Paixao *et al.* 2011). For each allele, the SD pattern is very stable (Phadke *et al.* 2014), but can be affected by environmental conditions such as temperature and nutrition during sexual reproduction. The probabilistic SD was previously shown to cause the evolution of uneven sex ratios in natural populations of *T. thermophila* (Paixao *et al.* 2011).

Like most ciliates, *T. thermophila* are facultatively sexual: cells reproduce asexually by binary fission when food is abundant and conjugation, the non-reproductive sexual stage, is induced between cells of different sexes under starvation conditions (Orias *et al.* 2011) (Figure S1 in File S1). Each *T. thermophila* cell contains a diploid germinal micronucleus (MIC) and a polyploid somatic macronucleus (MAC). During conjugation, the MIC undergoes meiosis to form gamete nuclei that fuse to produce new zygotic MIC via reciprocal fertilization. The new MAC differentiates from mitotic copy of the new MIC and goes through developmental genome editing and polyploidization, and the old MAC is destroyed by programmed nuclear death. A recent study revealed that the B-type *mat* allele in the MIC contained six pairs of incomplete genes, specifying sex II-VII, respectively (Cervantes *et al.* 2013). During MAC development, all but one gene pair is deleted and the remaining pair are re-assembled at the MAC *mat* locus by joining to intact transmembrane (TM) exons that are shared across all sex gene pairs. After conjugation, progeny will go through a period of sexual immaturity, typically lasting for 40–80 asexual generations (Perlman 1973), during which time cells are unable to mate. Knowledge about the patterns of probabilistic SD and molecular characterization of the *mat* allele makes *T. thermophila* a good multi-sex system in which to analyze the dynamics of sex ratio, from the perspective of population genetics.

Here, we first develop a population genetics model to dissect the mating kinetics in a large population of *T. thermophila* and to make quantitative predictions about how population sex ratio will evolve. We then investigate the impact of different parameters in the model on the dynamics of sex ratios. To test if population sex ratios follow the trajectories predicted by the model, we establish five replicate experimental populations and allow them to mate every 100 asexual generations, during which sex ratio dynamics and the frequencies of newly arisen mutations are tracked by time-course whole-genome sequencing. Overall, experimental observations highly support theoretical predictions. The newly established theoretical framework is applicable in principle to other multi-sex organisms and will provide more insight into the understanding of the maintenance of multiple sexes in natural populations.

## Materials and Methods

### The theory

Consider a large population of *T. thermophila*, in which sexual reproduction occurs periodically (for example, every 100 cell divisions) and in-between the population grows asexually.

Sexual reproduction starts with cell pairing (conjugation) which is assumed to be a random but synchronized process such that incompatible pairs (cells of the same sex) will dissolve and retry until no further pairing is possible. The process can be specified in detail as follows. Let *p*_*i*_ be the frequency of sex *i* right before the sexual reproduction and *q*_*i*_(*t*) be the relative frequency of sex *i* right after the *t*-th round of pairing, then *q*_*i*_(0) = *p*_*i*_. Since pairing is at random, at the *t*-th round of pairing the probability of a cell of sex *i* not paired is 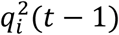, and after the frequency is re-calibrated, the frequency of sex *i* among the unpaired cells is

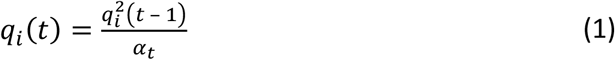

where 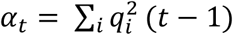, which is the expected relative proportion of unpaired cells after the t-th round of pairing. The overall proportion of unpaired cells will converge to

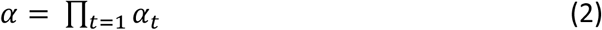

which is necessarily all cells of the most frequent sex type (*k*) before the sexual reproduction starts. Table 1 presents a summary of this and other parameters used in the description of the sexual reproduction process. After recalibration, the percentage of sex *i* among the cells involved in the pairing is

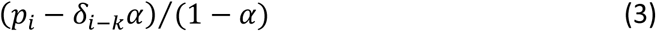

where *δ*_*m*_ takes value 1 if *m* = 0, and 0 otherwise.

In general not all paired cells proceed to the next step, some will dissolve and do not participate in the subsequent steps of the sexual reproduction together with the *α* proportion that are not paired. Let *β* represent the probability that each pair dissolves and furthermore it is assumed that each intact pair has probability *γ* to produce offspring (and has probability 1− *γ* to die without producing offspring). For each successfully reproducing pair, its two offspring have sex frequencies specified by the SD pattern *f* = (*f*_1_,…,*f*_7_). After the sexual reproduction, the total number of cells in the population is *Nr* where

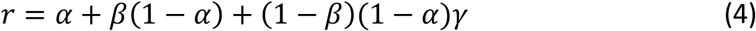

which is the probability that a cell either survives without participating in sexual reproduction or passes its genetics to offspring in sexual reproduction. Among the *Nr* cells, the proportion that is not sexual offspring is

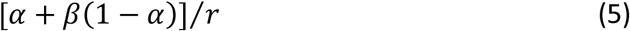

and there are

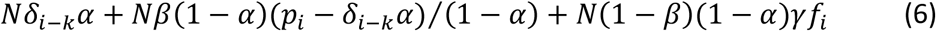

cells that are of sex *i*. Let 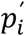 be the frequency of sex *i* after the sexual reproduction. It follows that

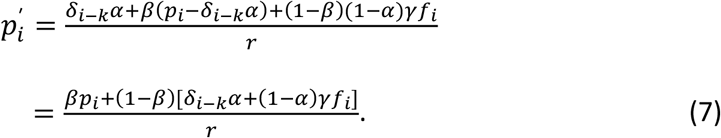

Since it is assumed that *N* is sufficiently large so that random genetic drift is negligible, 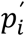 will be the frequency of sex *i* right before the next round of sexual reproduction if there is no natural selection.

By setting 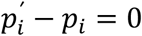, one can solve the equation for the equilibrium frequencies which lead to

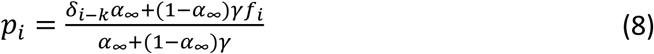

where *α*_*∞*_ is the limiting *α* value. It may appear that the equilibrium frequencies have nothing to do with *β*, but it is not true since its impact is reflected through the limiting *α* value.

Next we consider the impact of selectively advantageous mutations. Suppose a mutation emerges right after sex on a macronucleus of sex *i* with frequency 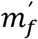 and growth rate 1 + *s* per cell division. Then right before the next sexual reproduction, the proportions of cells of various sexes (*j* = 1,…,7) are

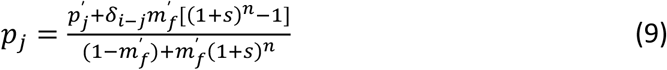

The frequency of cells containing the mutant right before sexual reproduction is

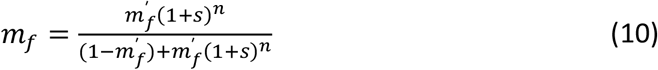

Immediately after the next sexual reproduction, mutant allele frequency becomes

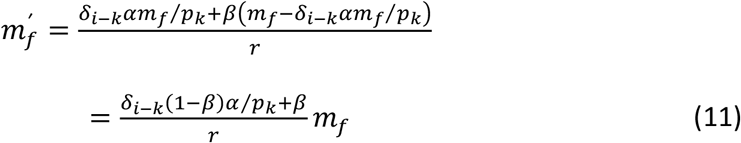

where *δ*_*i−k*_ is 1 if the mutant (sex *i*) is the most frequent sex and 0 otherwise.

By substituting the *m*_*f*_ in Equation (11) by (10), it leads to

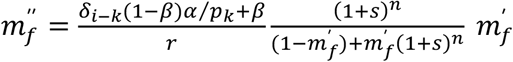

which leads to the equilibrium solution for *s* > 0

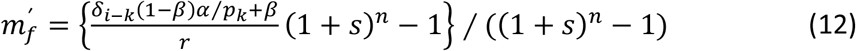

Apparently if *α* = 1, it will lead to the fixation of the mutant allele. Otherwise an internal equilibrium is possible.

If an advantageous mutation (in terms of growth) occurs in a micronuclear, and since the micronuclear is not expressed during growth, its advantage is hidden until it is passed to the macronuclear during sexual reproduction. The difference here is that the advantage is inheritable. Due to redistribution of sex, the mutant will spread over all sexes and eventually will be fixed without the necessity of fixing a sex.

### The experiment

#### Design of the experiment

To develop a better understanding on how various factors may affect the fate of a population, and to test if sex ratio dynamics follows our theoretical predictions, five replicate experimental populations (CS1-CS5) were established, each started with induced mating between equal proportion of homogeneous ancestor cells of sex IV and VI. The resulting population grew asexually for 100 generations with stable but large population size achieved by daily serial transfer and then it went into the next round of sexual reproduction. This sex:asex cycle was repeated 10 times for each of the replicate experiments. Immediately before each starvation to induce sexual reproduction, a large sample of cells was extracted, of which a portion was stored in liquid nitrogen (Cassidy-Hanley 2012) and the remaining was used for whole-genome sequencing for mutation identification as well as the determination of sex ratio. Due to the high cost of the whole-genome sequencing, we only sequenced DNA samples from CS1-CS3 across all time points and CS4 from generation 400 to 1,000. Although samples from CS5 were not sequenced, it still provided useful information on what sex was eventually fixed in the population. The essentials of the experiment are described below and more laboratory details can be found in the supplementary material (File S2).

#### Ancestral population

Strain SB210 mated with the star strain B*VII through genomic exclusion (GE) crosses (Allen 1967). The progeny of the GE crosses are whole-genome homozygotes in both nuclear genomes, but different progeny cells can exhibit different sexes in the MAC due to the probabilistic SD. When the progeny population matured, two cells, one of sex IV and another of sex VI (named Anc_IV and Anc_VI, respectively), were selected as ancestral cells and their asexual clonal populations became the starting cells for each of the replicate experimental populations. All these cells inherited the same B-type *mat* allele from SB210.

#### The parameters in the experimental populations

To estimate the parameters *β* and *γ*, only the proportional survival cells (*a*) and within which the proportion of asexual offspring (*b*) are required (when *α* is known). For example when *α* = 0, it follows from (4) and (5) that *β* = a × b and *γ* = *β* × (1 − *b*)/[*b* × (1 − *β*)]. The sex ratio pattern of the evolving population at the end of the first sex:asex cycle (i.e. at generation 100) was used to estimate *f*. As unmated Anc_IV and Anc_VI cells underwent serial passage along with progeny produced by sexual reproduction, correction for the background sex ratio of unmated cells was necessary to obtain the true *f* value representing the sex ratio pattern of sexual progeny. In addition, because natural selection during serial passage will also affect the estimation of *f*, we therefore recorded the daily growth rate as 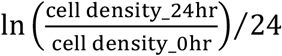 (Kishimoto *et al.* 2010) of the two ancestral populations in a week before mating and the resulting population during the following 100 asexual generations. For each evolving population, we tracked the changes in fitness by measuring the growth rate and determined parameter s as the ratio (minus 1) of the eventual population fitness and the starting ancestral population fitness that was calculated as the mean growth rate of the two ancestral populations over the week of serial passage. The relative fitness trajectory of each evolving population was analyzed by the SSlogis model in R 3.4.1 (http://www.r-project.org/).

#### Determination of sex ratio and identification of sexual progeny

The sex ratios in each sample were determined by mapping sequencing reads to the MIC reference genome (Hamilton *et al.* 2016). After removing the internally eliminated sequences (IESs) background within the *mat* locus, the sequencing depth of gene segments specific to each sex was used to determine the proportion of each sex. Two different approaches were used to determine whether unmated ancestral cells were responsible for the fixed sex. First, we performed sex testing experiments for all evolving populations at generation 600. Each evolving population was mixed separately with cultures of six test strains that are of sex II—VII, respectively (Hamilton and Orias 2000) to determine its fixed sex. Second, we used SNPs in the TM and truncated transmembrane (tm) exons of the MIC *mat* locus as molecular makers to distinguish sexual progeny from unmated ancestral cells because sexual progeny can acquire novel combinations of SNPs into MAC TM exons during mating (see details in File S2).

#### Whole genome sequencing, SNP calling and annotation

To track sex ratio dynamics and analyze the SNP in MAC TM exons to distinguish between sexual progeny and unmated ancestral cells, DNA samples were taken from populations CS1, CS2 and CS3 every 100 asexual generations until generation 1000. For SNP analysis in MAC TM exons in the population CS4, DNA samples were taken every 100 asexual generations from generation 400 to 1000. To prevent possible changes to the genetic structure of populations resulting from long-term storage, all DNA samples were isolated as soon as taken from each evolving population. However, due to the poor quality of the DNA sample from the CS2 population at generation 200, cell stock in liquid nitrogen was thawed and used for DNA isolation. Sequencing libraries were constructed using a standard Illumina protocol as previously described (Xiong *et al.* 2015). All libraries were sequenced to depths of about 30-fold coverage except libraries of the two ancestral samples that were sequenced to ˜250-fold coverage, using Illumina HiSeq 2000 or HiSeq 2500 instruments. A detailed SNP calling pipeline is given in File S2. Briefly, sequenced reads after trimming off adapters were aligned to *T. thermophila* MAC reference genome (Stover *et al.* 2006). Then we marked PCR duplicate reads, performed local realignment around potential indels and recalibrated the base quality score. We next ran VarScan 2.3.9 (Koboldt *et al.* 2009; Koboldt *et al.* 2012) for SNP calling. To reduce the risk of false positives, each mutation had to be supported by at least three forward and three reverse reads as previously reported (Sung *et al.* 2012; Long *et al.* 2016). By taking advantage of our time-course sequencing, we further refined candidate mutations based on mutation frequency trajectories, as previously reported (Lang *et al.* 2013; McDonald *et al.* 2016). In particular, the imprecise IES excision in the newly developing MAC during mating can result in the formation of many SNPs or indels around the IES junction sites (Hamilton *et al.* 2016). Therefore, candidate mutations were required to be located at least 60bp from the IES excision endpoint. Functional annotation of each mutation was carried out using SnpEff (Cingolani *et al.* 2012).

#### Identification and functional validation of beneficial mutations

We used two criteria to identify putative beneficial mutations. First, beneficial mutations are more likely than neutral or deleterious mutations to spread within a population. Thus, we considered mutations that reached a frequency of at least 0.9 as candidate beneficial mutations. Second, the frequency of beneficial mutations should correlate with changes in fitness. We therefore determined the Pearson correlation coefficients between changes in fitness and the frequency of each identified mutation. Putative beneficial mutations were required to have a correlation coefficient of ≥+0.8 (Table S2). Mutations that met both criteria were classified as beneficial. To validate whether a selected mutation confers a fitness benefit, we introduced it into both ancestral cells, see supplementary material in File S2. Then we mixed each mutant cell population with its corresponding ancestral cell population in roughly equal proportion and propagated them under the same conditions as the evolution experiment. Sanger sequencing was performed every two days to determine the relative proportion of each cell type. PCR primers used to amplify sequences contain mutation site were: Mut-f4001 5′-TAGATTAAGACACTTTAGAAAAAGC-3′ and Mut-r4898 5′-TCATTGATTCATTAGATTATCTTTC-3′. Competition assays were carried out in duplicate.

## Data availability

The mathematic model was analyzed using Java 9 and the source code will be available upon request. File S1 contains additional figures including the life cycle of *T. thermophila* (Figure S1), impact of the model parameters on sex ratio patterns when using another *f* (Figure S2), and growth rate records of the two ancestral populations and the three sequenced evolving population during the first 100 asexual generations (Figure S3). Table S1 in File S1 shows the sex ratios in the three sequenced evolving populations at generation 100. File S2 contains additional Materials and Method sections. Table S2 contains frequency trajectories and functional annotation of all identified mutations. Supplementary material has been uploaded to figshare. All genome sequencing data are available at Sequence Read Archive with accession number SRP080979.

## Numerical Results

### Impact of the model parameters on the equilibrium frequencies of sexes

As shown in the Theory section, parameters influencing the equilibrium sex ratios include the initial population sex ratio before mating, *β*, *γ*, *f* and *s*. We thus explored numerically the impact of these parameters.

Define *f*_*max*_ as the largest *f*_*i*_ and *sf*_*max*_ as the corresponding sex. To simplify the illustration, we first set *f* to be the estimated SD pattern (II, 0.306; III, 0.010; IV, 0.267; V, 0.047; VI, 0.113; VII, 0.257) from our experiments. Then, *f*_*max*_ = 0.306 and *sf*_*max*_ = II. We started by considering the scenario in which the population starts with two randomly selected sexes of equal proportions (Figure 1, A-D). Figure 1A shows that the population sex ratios at equilibrium exhibit two patterns under different combinations of *β* and *γ*: all six sexes co-exist or a single sex is fixed. Specifically, larger *γ* facilitates the co-existence, and the probabilities of fixing different sexes are positively correlated with the sex frequency distribution in *f*. In addition, Figure 1, B-D indicates larger *β* can increase the probability of fixing other sexes except the *sf*_*max*_. We then allowed in the initial population random sex ratio of the given set of sexes (but their frequencies sum necessarily to 1). Figure 1E shows that random sex ratio remarkably decreases the fixation probability of *f*_*max*_ compared to that shown in Figure 1A. However, increasing the number of sexes in the initial population promotes the fixation of *f*_*max*_ (Figure 1F). These comparisons show that higher sex diversity, including more even sex ratio and greater number of sexes, facilitates the fixation of *f*_*max*_. In the extreme case, when a population starts with equal proportions of all the six sexes, only the *sf*_*max*_ (sex II) can be fixed (Figure 1G). Similar results were also found when *f* is set to other SD patterns, such as the one described in the Introduction section (Figure S2 in File S1).

**Figure 1.**
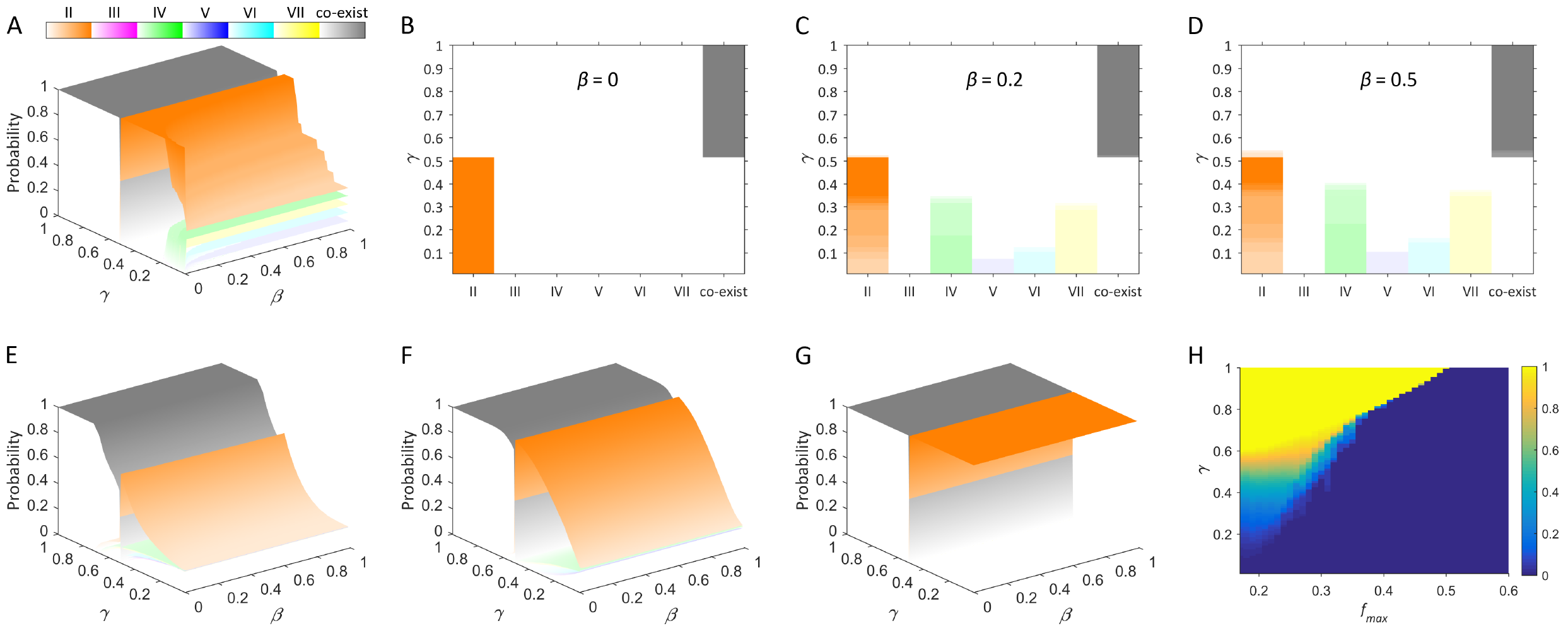
Impact of the model parameters on sex ratio patterns when there is no selection. (A-D) Sex ratio pattern at equilibrium when the population starts with equal proportions of any two of the six sexes (A) and under three specific *β* values (B-D). The gray color represents the situation that all sexes co-exist and the other six colors represent the fixation of one of the six sexes II-VII, respectively. The color gradients represent the probability of occurrence for each situation. This color scheme applies to all panels except panel H. (E,F) Sex ratio patterns at equilibrium when the population starts with any two of the six sexes (E) or all the six sexes (F) and the initial sex ratio is random but the sum is 1. (G) A special case of sex ratio pattern when the population starts with even proportions of six sexes. (H) The probability of co-existence of the six sexes under different values of *f*_*max*_, the most frequent sex in parameter *f* and *β* is set to 0.

Next, we evaluated the impact of *f*_*max*_ values on the patterns of equilibrium sex ratios. For a B-type *mat* allele, the *f*_*max*_ value can range from 1/6 to 1. Figure 1H shows the probability of co-existence of all sexes decreases with the increase of *f*_*max*_. In particular, when *f*_*max*_ is greater than 0.5, co-existence will not occur. A simple explanation is that when more than half of the progeny produced by any two of the six sexes in a random-mating population express the sex of *f*_*max*_, the frequency of *f*_*max*_ will gradually increase up to fixation.

We further investigated how mutation that occurs on the MAC right after sexual reproduction affects the evolutionary consequence of sex ratios. For simplicity, we only considered the two cases in which either the *sf*_*max*_ (sex II, Figure 2A) or the least frequent one (sex III, Figure 2B) obtains a beneficial mutation, and assumed that the population starts with equal proportions of any two of the six sexes. Compared to Figure 1A, Figure 2 shows that the patterns of equilibrium sex ratio changes strikingly. Sex obtaining a beneficial mutation has an increased probability to be fixed and larger *β* or *s* values will intensify this tendency. When *s* is larger than a certain threshold, sex with the beneficial mutation will be destined for fixation regardless of the values of *β* and *γ*.

**Figure 2.**
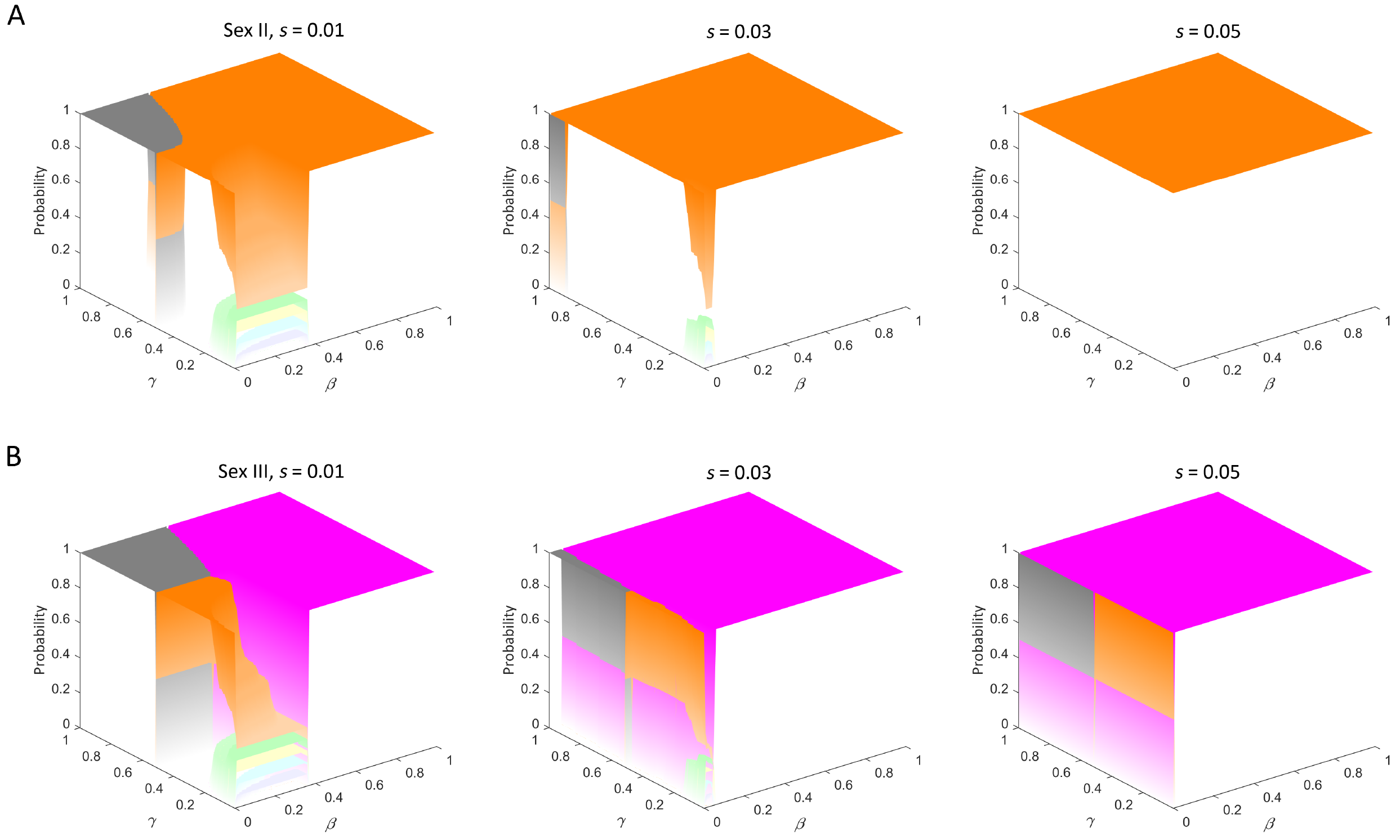
Impact of selection on sex ratio patterns. Sex ratio pattern at equilibrium when sex II, the most frequent sex. (A) and sex III, the least frequent sex (B) in parameter *f* obtain a beneficial mutation with different selective coefficients. Color regime is the same as Figure 1A.

### Impact of environmental fluctuations on the equilibrium frequencies of the sexes

In the numerical exploration described above, *f* is fixed which is relatively stable for a given experimental setting. However, in reality change in some environmental factors may lead to the modification in the values of *f* and other parameters. In natural populations of *T. thermophila*, a series of environmental fluctuations including the changes of abiotic and biotic factors are possible to influence the equilibrium sex ratio pattern that has already established in the ideal mating population. For example, migration among populations leads to the immediate changes of population sex ratio, temperature and food availability affect the parameter *f* (Nanney 1960; Orias and Baum 1984) and a salty environment can changes the values of *β* and *γ*.

For simplicity of illustration, consider a population in which cells will not dissolve once successfully paired together (*β* = 0) and all pairs will produce progeny (*γ* = 1), its sex ratio at equilibrium can be readily predicted when the parameter *f* is determined. For example, a population with the empirically estimated *f* will reach the following equilibrium sex ratio: II, 0.316; III, 0.010; IV, 0.263; V, 0.046; VI, 0.112; VII, 0.253. By imposing a varying degree of deviations or fluctuations on either the established equilibrium sex ratio (Figure 3A) or the parameter *f* when determining the progeny sex distribution after each round of mating (Figure 3B), we found sex ratio patterns are both changed dramatically and the impact of fluctuation on parameter *f* seems more remarkable. A slight fluctuation will lead first to the fixation of the *f*_*max*_, and with the increase of fluctuation degrees, other sexes can also be fixed with probabilities positively correlated with the sex frequency distribution in *f*.

**Figure 3.**
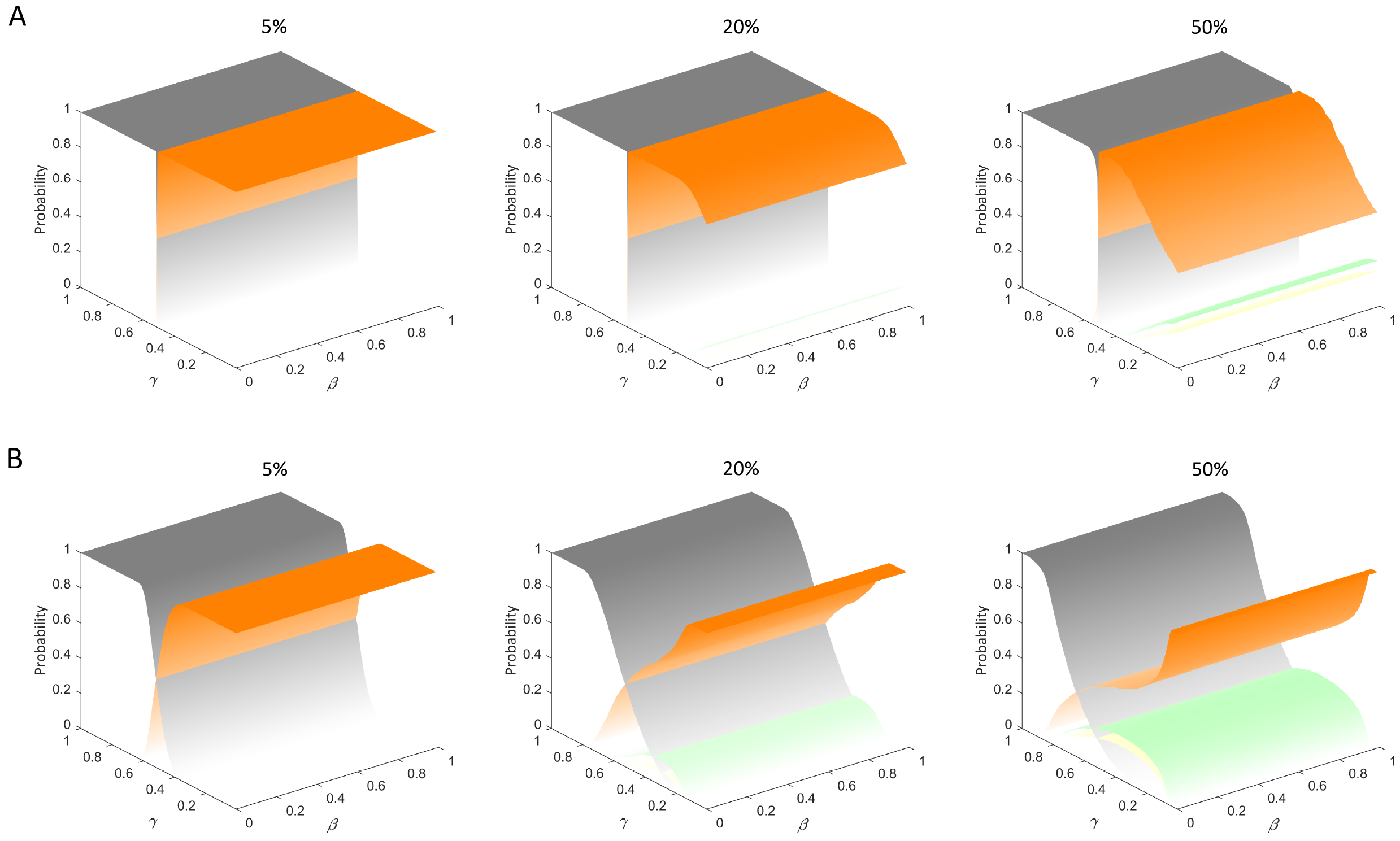
Impact of environmental fluctuations on sex ratio patterns. Sex ratio pattern at equilibrium when the established equilibrium sex ratio. (A) or the empirically estimated parameter *f* (B) fluctuates with varying degrees. Color regime is the same as Figure 1A.

## Experimental results

### Parameter estimation and prediction of sex ratio dynamics

After mating between the two ancestral populations of equal proportion (*α* = 0), we found in the five replicate experimental populations, the number of cells decreased to an average of 52% within which 38% were the two unmated ancestral cells and each of them should account for 19%. According to equation (4) and (5), parameter *β* is equal to 20%, and *γ* is equal to 40%. As there was no significant difference in growth rate among the two ancestral populations and the resulting population after mating (Figure S3 in File S1), it is reasonable to assume that the proportion of each unmated ancestral cell (19%) did not change significantly and the effect of natural selection could be neglected during serial passage for the first 100 generations. Thus, after correction by subtracting background sex ratio of unmated ancestral cells from the average sex ratio (Table S1 in File S1) in the three sequenced evolving populations (CS1-CS3) at generation 100, the *f* values were: II, 0.306; III, 0.010; IV, 0.267; V, 0.047; VI, 0.113; VII, 0.257.

Using these estimated parameter values, we could readily predict the sex ratio dynamics in a large experimental population. When a population begins with the two ancestral populations in equal proportions and mates every 100 asexual generations, and there is no selection, we found sex II would increase in frequency gradually and eventually be fixed (Figure 4B).

**Figure 4.**
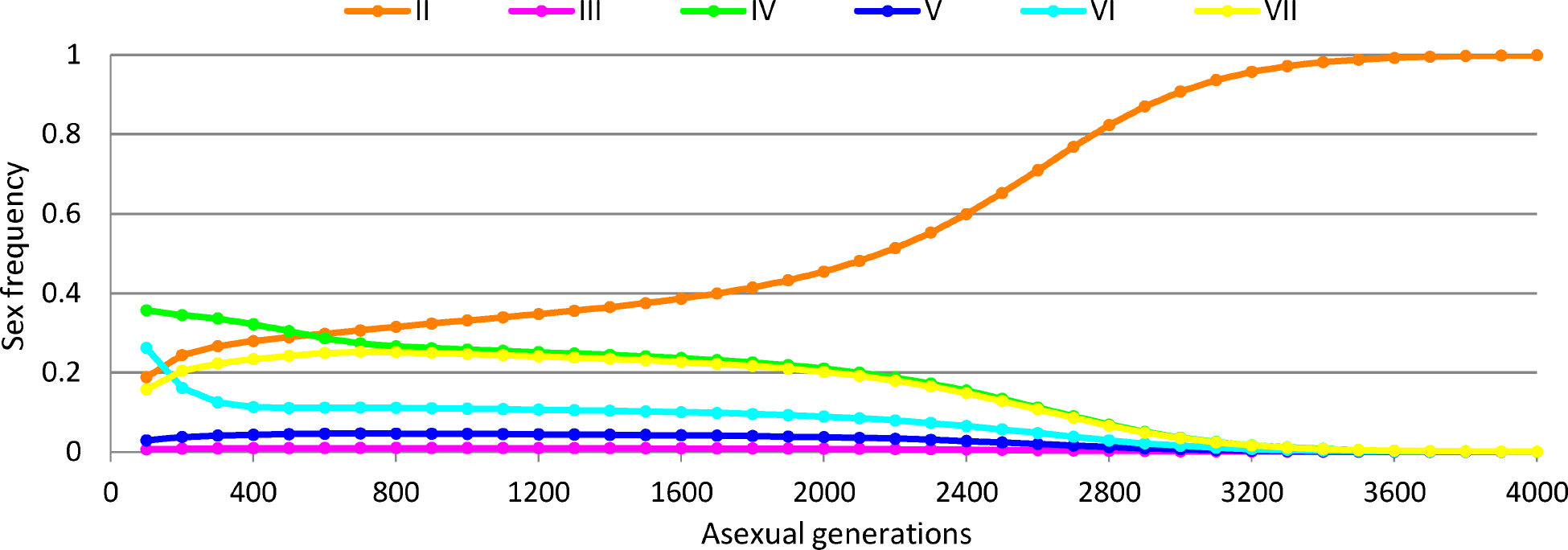
Prediction of sex ratio dynamics. Predicted sex ratio trajectory when populations start with equal proportions of the two ancestral populations and mate every 100 asexual generations, and there is no selection.

### Natural selection is responsible for the fixation of single sex

Sex ratio dynamics showed that at generation 100, all three sequenced populations (CS1-CS3) contained a mixture of cells with sex II-VII. Over time, however, a different single sex became dominant gradually and then fixed within each population (Figure 5A). In addition, during each round of mating, we calculated the ratios of mating pairs and sexual progeny (File S2). After several sex:asex cycles, no mating pairs or sexual progeny were observed in any of the replicate populations (Table 2). Sex testing at generation 600 showed that each population expressed only a single sex, but there were four different sexes fixed in the five replicate populations. These results deviated strikingly from the prediction that only sex II could be fixed if all parameters were the same and there is no selection. Considering the large population size in our experiment, the observed deviations are unlikely to be caused by genetic drift. A more plausible explanation is that a subgroup of cells expressing a specific sex gain an increased growth fitness and expanded to fixation consequently. Indeed, the overall growth record revealed that the relative fitness of each evolving population increased markedly (Figure 5B).

**Figure 5.**
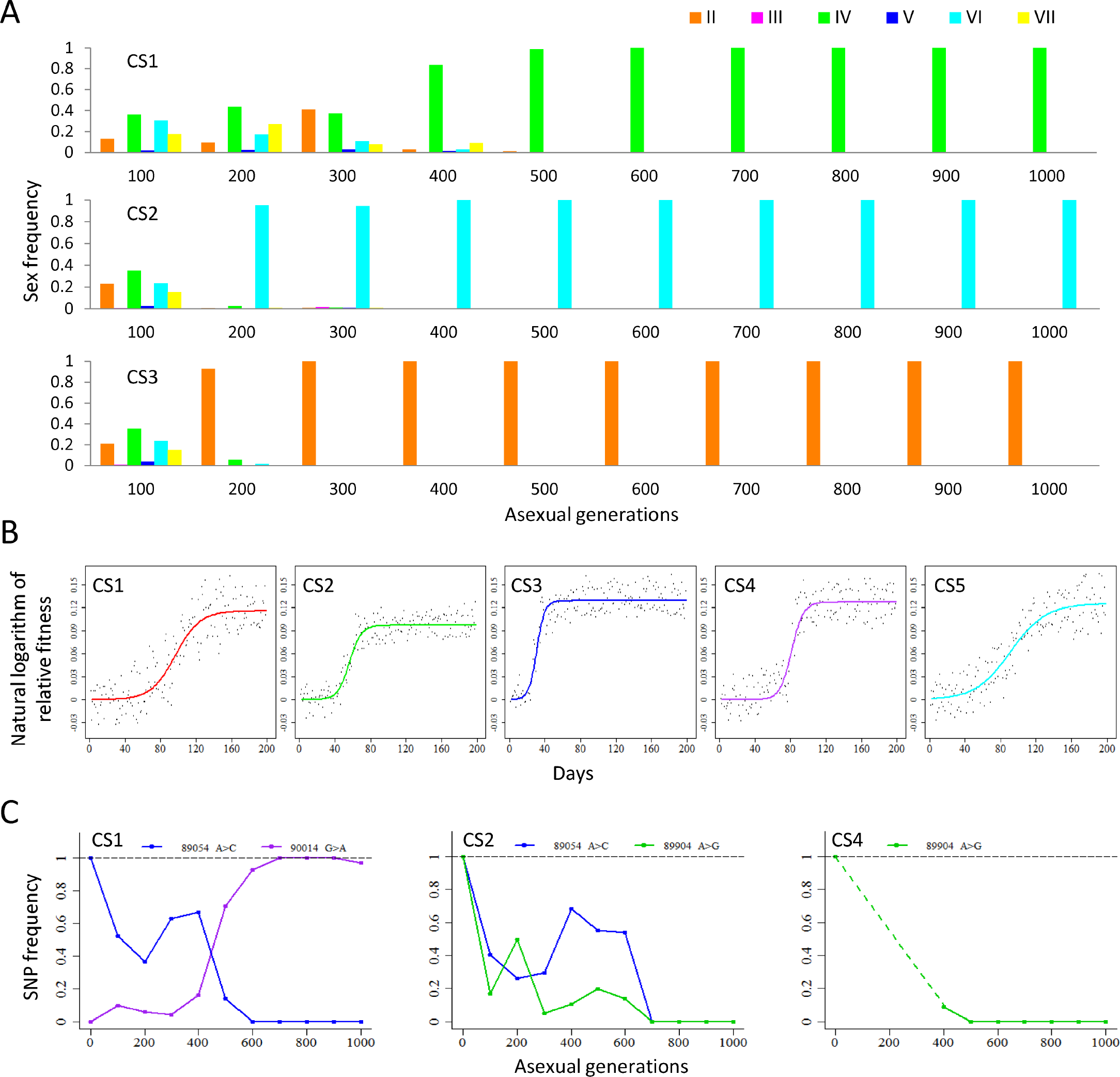
Increased fitness of sexual progeny were responsible for the fixation of a certain sex in each of the replicate populations. (A) Sex ratio dynamics in the populations CS1 to CS3. Each colored bar represents the frequency of a sex, as determined by sequencing data. (B) Relative fitness trajectories during experimental evolution. The fitted growth model for each replicate population was plotted with a different colored line. (C) Temporal changes in the frequency of novel SNPs recombined into progeny MAC TM exons. Panels show SNP frequency changes in the CS1, CS2, and CS4 populations. Numbers indicate SNP positions at the MIC *mat* locus and frequency changes refer to non-reference bases, e.g. in the CS1 panel, for A > C, A is the reference base and C is the non-reference base. Generation 0 represents SNP frequency in the two ancestral cells. Each SNP is represented by the same color in all three panels.

In three of the replicate populations (CS1, CS2, and CS4), the fixed sexes were the same as one of the ancestral cells, sex IV or VI. However, according to SNP analysis in the MAC TM exons, we identified several SNPs that were either lost or became fixed in each of the three populations (Figure 5C). In addition, the other two populations, CS3 and CS5, fixed different sexes (II and VII, respectively) than the ancestral cells. Thus, unmated ancestral cells had been completely replaced in all the five replicate populations, suggesting that sexual progeny cells might gain a faster growth rate than ancestral cells.

### Identification of beneficial mutations

The analysis described in the previous section establishes that increased fitness of sexual progeny was responsible for the fixation of a certain sex in each of the replicate populations. To reveal the molecular basis underlying the increased fitness, we turn to whole-genome sequencing data. We detected an average of 14.7 *de novo* mutations per population and several candidate beneficial mutations were identified in the three sequenced populations: two in CS1, one in CS2, and one in CS3. We then investigated whether a selected candidate beneficial mutation from the CS1 population confers a fitness benefit (Figure 6A; red line). This point mutation resulted in a premature termination codon in a gene encoding a serine/threonine kinase (Table S2). Within 12 days, ancestral cells had been completely replaced by mutant cells (Figure 6B), providing clear evidence that the selected mutation conferred increased growth fitness.

**Figure 6.**
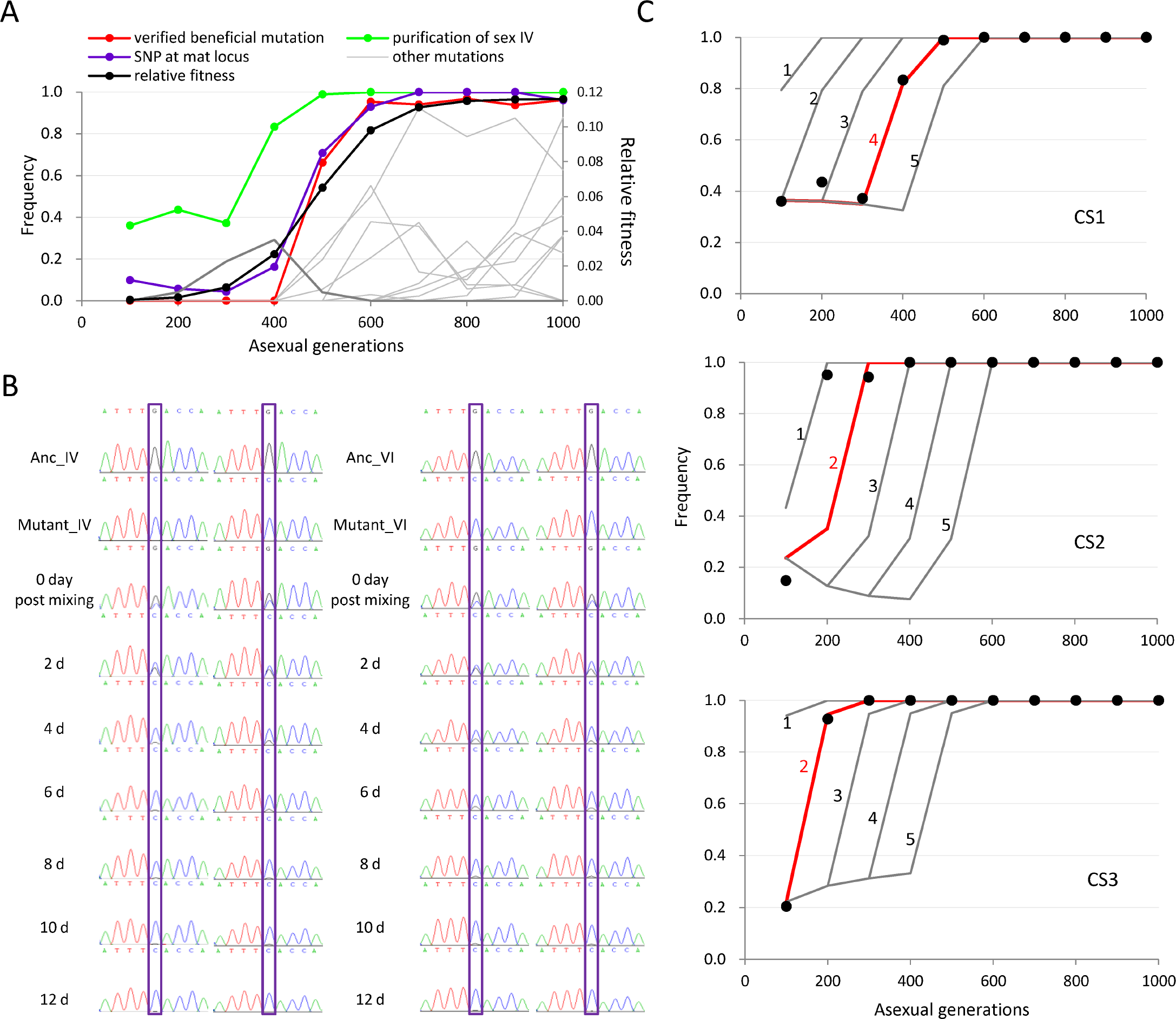
The fixation of a single sex in the population is caused by newly arisen beneficial mutations. (A) Interactions between beneficial mutations and sex fixation in the CS1 population. Red line, a verified beneficial mutation; gray lines, other detected mutations. The green shows the process of sex IV purification within the population. The black line shows the growth fitness trajectory. The purple line represents the change in frequency of the fixed novel SNP at the MAC *mat* locus (shown in Figure 5C; purple line in CS1). (B) Functional validation of putative beneficial mutations. Each of ancestral cell populations was co-cultured with its corresponding mutant cell population containing the candidate beneficial mutation (panel A; red line) with daily serial passage. DNA samples were taken every 2 days and analyzed by Sanger sequencing to determine the relative ratio of the two cell types. Competition assays were performed in duplicate. (C) Comparison of sex fixation trajectories between experimental observations and the model’s predictions. Model predictions assume the fixed sex in each population acquired a beneficial mutation in the MAC after one of the first five rounds of mating. Black dots are observed frequencies. The best fits are shown as red lines.

Given that genes determining sexes are not expressed during growth, it seems unlikely that they can confer a selective advantage leading to their own rapid fixation. However, in the case of the CS1 population, purification of sex IV preceded the fixation of all candidate beneficial mutations. This apparent paradox might be explained in the following way: directly after a single sex becomes purified, the population might still contain two different cell types of the same sex, with only one of these containing a beneficial mutation. In the CS1 population, sex IV had become purified by about generation 500 (Figure 6A; green line). However, the novel SNP at MAC *mat* locus, a marker for the eventually fixed sex IV genes, did not become fixed until about generation 600 (Figure 6A; purple line). This suggested that not all MAC *mat* loci exhibiting sex IV contained this novel SNP at generation 500, and that over the following 100 asexual generations (from generation 500 to 600), those sex IV cells containing the novel SNP swept rapidly to fixation by outcompeting sex IV cells lacking the novel SNP, a phenomenon known as “clonal interference”. Notably, frequency changes in the novel SNP and the verified beneficial mutation correlated strongly (*r* = 0.99). This result suggested that sex IV cells containing the novel SNP also acquired this verified beneficial mutation, and sex IV genes within the cells became fixed through genetic hitchhiking with the beneficial mutation.

Moreover, the effect of selection due to newly arisen beneficial mutations was likely to be the cause that trajectories of sex ratio in experimental populations deviated strikingly from theoretical prediction that assumed no selection. According to the fitness trajectories, the values of s in three sequenced populations are: 0.12 in CS1, 0.10 in CS2, 0.14 in CS3. For each population, we then assumed a mutated cell expressing the fixed sex acquired the corresponding selective advantage in the MAC right after one of the first five rounds of mating, respectively. In fact, we did not detect the verified beneficial mutation of the CS1 population in the MIC sequencing data (data not shown). The results showed the predicted frequency trajectory of the fixed sex could highly agree with the observation for each population under a specific assumption (Figure 6C). The relative low agreement in the CS2 population might have resulted from a DNA sample at generation 200 being isolated from the thawed cultures stored in liquid nitrogen, and the long-term storage might change the composition of population sex ratio. Overall, when considering the effect of selection, the results showed that the experimental observations highly supported our model’s predictions and newly arisen beneficial mutations were the cause of the fixation of single sexes in the populations. In addition, the agreement between experimental observations and model predictions in the three sequenced populations (CS1-CS3) suggested that the fixation of a single sex in the populations CS4 and CS5 was also probably due to newly arisen beneficial mutations because both of these two populations showed a remarkably increased growth fitness (Figure 5B) and fixed a different sex (IV and VII, respectively) than sex II that is predicted to be the only fixed sex when there is no selection (Figure 4).

## Discussion

In organisms with two sexes, male and female, the strategy to adjust population sex ratio has been well demonstrated. Nonetheless, how organisms with multiple sexes maintain proper sex ratio in the population remains poorly understood. Based on a newly developed population genetics model, we analyzed in this study the dynamics of population sex ratio in *T. thermophila*, a ciliate with seven sexes and probabilistic SD mechanism. We found there are plenty opportunities for both the co-existence of all sexes and the fixation of a single sex, depending on the combinations of several parameters, including the strength of natural selection. Specifically, parameter *γ* mainly determines which pattern the population will exhibit, and the probability that a specific sex is fixed is positively correlated with its frequency in parameter *f*, but it is also affected by *β* (Figure 1, B-D). Natural selection can strongly shift the sex ratio pattern toward the fixation of a single sex with a growth advantage. Moreover, in natural populations of *T. thermophila*, all of these parameters likely fluctuate along with changing environmental conditions, which can further diversify the population sex ratio patterns (Figure 3). Experimental observations of sex ratio dynamics confirmed the validity of the model’s predictions.

In natural habitats of *T. thermophila*, all seven sexes were generally present (Doerder *et al.* 1995). However, sex ratios varied remarkably in different sampled ponds and even displayed local and seasonal variation in the same pond. It was postulated that sex ratio variations between and within ponds are due to the fluctuations of SD pattern through the interaction between multiple *mat* alleles and environmental conditions (Arslanyolu and Doerder 2000). This hypothesis is consistent with our model prediction that there is sufficient opportunities for the coexistence of all seven sexes and environmental fluctuations can further increase the diversity of sex ratios. Moreover, it suggests that population sex ratios at equilibrium are determined by not only the SD pattern *f* but also other parameters involved in sexual reproduction (equation 9). Thus, our theoretical model provides a comprehensive framework to illustrate how natural populations of *T. thermophila* maintain proper sex ratios when responding to changeable environments.

Our numerical exploration also suggests that there is a large probability of fixation of a single sex in a local population. Indeed, due to limited rates of dispersal in *T. thermophila* (Zufall *et al.* 2013), local populations or subpopulations sampled from the same pond often contained only a single sex even though all sexes are present in the pond (Doerder *et al.* 1995). The fixation of a single sex in a location can potentially provide an advantage to prevent inbreeding because a particular consequence of probabilistic SD is that it allows for mating among genetically identical individuals at the micronucleus. The experimental results combined with the model’s predictions suggest that the fixation of a single sex is often caused by beneficial mutations during sexual reproduction. Interestingly, the process of choosing which sex to be fixed seems to be random because four different sexes are fixed in the five replicate experimental populations. Thus, it is likely that in natural subpopulations, the spontaneous beneficial mutations help their carrying sex to fixation, which effectively blocks further local inbreeding, and then the subpopulations can expand rapidly through asexual production. However, due to the probabilistic SD, mating between subpopulations only containing two different sexes could lead to the recovery of all the seven sexes, which facilitates the population to regain the benefits provided by sex when the environment changes dramatically. Thus, the particular probabilistic SD may provide an advantageous mechanism for the lineage survival of *T. thermophila* by allowing this species to integrate the benefits of both sexual and asexual reproduction. However, why the number of sexes is fixed at seven in natural populations of *T. thermophila* remains a mystery. In fact, the *Tetrahymena* species have various number of sexes ranging from three to nine and display different modes of sexual inheritance (Phadke and Zufall 2009), but little is known about their SD mechanisms, which needs further sequencing of the MIC genome to elucidate. When more SD mechanisms and sex ratio data from experimental evolution and natural populations in different species are available, comparative analysis may be used to explore if there exists an optimal number of sex for a specific species to adjust sex ratio dynamics in the light of our theoretical framework.

In summary, our theoretical model combined with experimental results provides a comprehensive framework to analyze the dynamics of sex ratio in *T. thermophila*, and proposes a possible strategy of maintaining multiple sexes in natural populations of *T. thermophila*. In principle, the newly established theoretical framework is applicable to other multi-sex organisms to bring additional insight into the understanding of maintenance of multiple sexes in a natural population.

## Acknowledgments

We thank Eduardo Orias (University of California Santa Barbara) and Yong Zhang (Institute of Zoology, Chinese Academy of Sciences) for helpful discussions during the study. Additional students in Miao’s laboratory provide assistance in the long-term cell culture: Wentao Yang, Guanxiong Yan, Zongyi Sun, and Chuanqi jiang. This study was supported by the National Natural Science Foundation of China (grant number 91631303 to W.M. and 91631304 to Y.X.F).

